# Genome-wide local ancestries discriminate homoploid hybrid speciation from secondary introgression in the red wolf (Canidae: *Canis rufus*)

**DOI:** 10.1101/2020.04.05.026716

**Authors:** Tyler K. Chafin, Marlis R. Douglas, Michael E. Douglas

**Author notes:** Email: (TKC), send reprint requests to this address; (MRD); (MED).

## Abstract

Hybridization is well recognized as a driver of speciation, yet it often remains difficult to parse phylogenomically in that post-speciation gene flow frequently supersedes an ancestral signal. Here we examined how interactions between recombination and gene flow shaped the phylogenomic landscape of red wolf to create non-random retention of introgressed ancestry. Our re-analyses of genomic data recapitulate fossil evidence by demonstrating red wolf was indeed extant and isolated prior to more recent admixture with other North American canids. Its more ancient divergence, now sequestered within low-recombinant regions on the X-chromosome (i.e., chromosomal ‘refugia’), is effectively masked by multiple, successive waves of secondary introgression that now dominate its autosomal ancestry. These interpretations are congruent with more theoretical explanations that describe the manner by which introgression can be localized within the genome through recombination and selection. They also tacitly support the large-X effect, i.e., the manner by which loci that contribute to reproductive isolation can be enriched on the X-chromosome. By contrast, similar, high recombinant regions were also found as enriched within very shallow gene trees, thus reflecting post-speciation gene flow and a compression of divergence estimates to 1/20^th^ of that found in recombination ‘cold spots’. Our results effectively reconcile conflicting hypotheses regarding the impact of hybridization on evolution of North American canids and support an emerging framework within which the analysis of a phylogenomic landscape structured by recombination can be used to successfully address the macroevolutionary implications of hybridization.

## Introduction

Hybridization was once considered a rare event. However, its adaptive potential as a macroevolutionary process (i.e., an unfettered access to an extensive panoply of genetic variation; Grant and Grant 2019) has been enhanced by the widespread adoption of genomic approaches (Abbott et al. 2013; Twyford and Ennos 2012; Taylor and Larson 2019). As such, hybridization has now become one component of a more contemporary approach to species diversification. Prior to the onset of genomics, there were few examples of homoploid hybrid speciation in animals (Mavarez and Linares 2008), with notable exceptions being the Virgin River chub (*Gila seminuda*; DeMarais et al. 1992; Chafin et al. 2019) and the red wolf (*Canis rufus*; Wayne and Jenks 1991; Reich et al. 1999). Being that hybrid speciation is now becoming a common hypothesis (Yakimowski and Rieseberg 2014; Elgvin et al. 2017; Lamichhaney et al. 2018; Eberlein et al. 2019; Marques et al. 2019), we argue that a framework must now be developed so as to discriminate among its alternative outcomes [e.g. a more explicit definition of ‘homoploid hybrid speciation;’ (Schumer et al. 2014; Nieto Feliner et al. 2017; Schumer et al. 2018)].

Given the inherent difficulties associated with diagnosing hybrid speciation as the basis of reproductive isolation, it has often been defined on the basis of genomic mosaicism (Blanckaert and Bank 2018; Schumer et al. 2018a). However, doing so risks overlooking a more varied evolutionary role for hybridization. A contributory aspect is the recognized difficulty in detecting hybridization, for it is but one of several mechanisms driving phylogenetic discordance in the genome (Maddison 1997; Degnan and Rosenberg 2009). Hybridization–speciation dynamics are further complicated by the fact that evidence of archaic branching (i.e. those that precede introgression) can be depleted, and especially so in those lineages with a history of secondary introgression. However, the parsing of genealogical histories is dependent on the interactions between recombination, genetic drift, and selection (McGaugh et al. 2012; Schumer et al. 2018). As such, branching patterns are often retained non-randomly, with reduced permeability to gene flow found in those genomic areas with low recombination, where introgression of deleterious alleles is restricted by an increased efficacy of linked selection (Payseur and Rieseberg 2016; Runemark et al. 2018; Schumer et al. 2018).

The interaction between selection and recombination through time allows fundamental predictions to be made with regard to the stability of hybrids genomes, and this may promote the role that hybridization plays in a given lineage. In the generations following a hybridization event, recombination creates junction-points where ancestries transition from one parental genome to another (Fisher 1954). Their densities along the length of a chromosome can be used to find loci relating to hybrid fitness, because selection against incompatible loci will alter the breadth of correlated ancestry, depressing local recombination with proportionately larger distances between junctions (Sedghifar et al. 2016; Hvala et al. 2018).

We thus hypothesized if signatures of archaic introgression are indeed masked by secondary introgression, then the probability of observing the ‘original’ ancestry will increase as local recombination rates decrease [even when hybrid ancestries dominate, as is sometimes the case (Fontaine et al. 2015)]. Thus, our prediction is that patterns of coalescence will be multimodal, reflecting the times and manner by which populations have diverged and subsequently intermingled (Rosenberg and Feldman 2002). Here we explore how this distribution in the red wolf is shaped by genome structure and recombination rate heterogeneity. To do so, we test multiple opposing hypotheses regarding the role hybridization has played in the history of this species.

### The red wolf as a case study

Our capacity to more precisely delineate hybridization has precipitated ancillary issues, such as the disparity that now exists between evolutionary complexity and species conservation (Ellstrand et al. 2010; Fitzpatrick et al. 2015; Supple and Shapiro 2018; vonHoldt et al. 2018). The U.S. Endangered Species Act (ESA 1973; 16 U.S.C. § 1531 et seq), as well as similar legislations globally, do not protect hybrids (Jackiw et al. 2015), despite scientific support (O’Brien and Mayr 1991; Allendorf et al. 2001; Haig and Allendorf 2006; Lind-Riehl et al. 2016). Few species have been as integral to this debate as red wolf (*Canis rufus*), fueled in part by the long-standing ambiguity surrounding its origins (Gittleman and Pimm 1991; Wayne and Jenks 1991; Dowling et al. 1992; Nowak 1992).

DNA evidence implicates hybridization, which some have attributed to recent coyote (*C. latrans*) and grey wolf (*C. lupus*) admixture (Wayne and Jenks 1991; Roy et al. 1996; Reich et al. 1999; vonHoldt et al. 2011; vonHoldt et al. 2016). Alternatively, others have instead argued that data point to an earlier red wolf origin, with introgression occurring as a subsequent phenomenon (Nowak 1979; Dowling et al. 1992; Nowak 1992; Wilson et al. 2000; Nowak 2002; Hohenlohe et al. 2017). Hypotheses regarding the status of red wolf are as follows (per Waples et al. 2018). It is: (1) An evolutionary distinct lineage derived from common ancestry with either *C. lupus* or *C. latrans* (=secondary introgression); (2) A transient product produced by contemporary hybridization (=hybrid swarm), or (3) An admixture subsidiary to a more ancient hybridization (=hybrid speciation). We discriminate among these scenarios by establishing predictions with regard to the respective footprint each would leave on the genomic landscape of red wolf, then testing each to ascertain which has the greatest probability of occurrence.

## Results

From prior publications (vonHoldt et al. 2016; Kukekova et al. 2018), we obtained whole-genome sequences for the red wolf (*Canis rufus*) and its putative progenitor species [the North American gray wolf (*C. lupus*) and coyote (*C. latrans*)], as well as an outgroup species (red fox; *Vulpes vulpes*). We aligned these data against the domestic dog genome (Kirkness et al. 2003; Lindblad-Toh et al. 2005; Hoeppner et al. 2014), then extracted full chromosome-length ‘pseudoalignments’ from all nucleotide positions having sufficient sequencing depth. This resulted in an average of 95.5% of the genome having called bases across species.

To identify sub-genomic ancestry blocks, we partitioned the 38 autosomes and the X chromosome into 913,849 non-overlapping windows by using an algorithm that defined a ‘most parsimonious’ set of hypothesized ancestry breakpoints, given a four-gamete assumption (Chafin 2020). This provided data with an average length of 2.2 kb (10.3 kb if merging consecutive ancestry blocks; see Fig. S1). We then analyzed each chromosome separately, and additionally partitioned regions by recombination rate, as inferred using an existing high-density linkage map (Fig. S2)(Wong and Neff 2009; Wong et al. 2010).

Our analyses are presented in two stages: The first examines the distribution of phylogenies across the genome. Here, we reasoned that sub-genomes from the putative parental lineages could be assigned via Maximum Likelihood (ML) estimates of the local branching order. We then identified heterozygous ancestry blocks by calculating the interspecific heterozygosity of red wolf sequence within each sub-alignment. We established relatively simple predictions for these early analyses: If indeed hybridization is recent (per Wayne and Jenks 1991; vonHoldt et al. 2016), then ancestry blocks will be large, given scant time for linkage blocks to be broken up by recombination (Falush et al. 2003; Pool and Nielsen 2009)]. Also, interspecific heterozygosity should be high (Rieseberg and Linder 1999; Anderson and Thompson 2002), whereas raw divergence will be low (vonHoldt et al. 2017). Likewise, we also expect an enrichment of introgressed histories in regions with higher recombination (Schumer et al. 2018; Li et al. 2019). We then used a model of linkage block decay to test several alternative models of hybridization, with gene flow either being gradual (e.g. declining through time from an initial event), continuous, or have occurred in multiple independent waves (Ni et al. 2018). However, when gene flow is high, signals of more ancient divergence could be ‘masked.’

To untangle this, our second approach employed local phylogenetic signals as latent variables within a hidden Markov model [=coalHMMs (Dutheil et al. 2009; Spence et al. 2018)]. It allowed us to extract parameter estimates (e.g. divergence times) by integrating results from coalescent theory, despite the fact that the ‘true’ history at each nucleotide is masked (Hobolth et al. 2007; Dutheil et al. 2009). We then contrasted this approach with a second coalescent-based method [g-PhoCS; (Gronau et al. 2011)] that replicates the analyses of vonHoldt et al. (2016) with the exception that inputs were additionally partitioned by their respective sub-genomic histories.

### Representation and divergence of parental genomes

Phylogenetic estimation and interspecific heterozygosity revealed that 26.8–36.5% of ancestral blocks were heterozygous (Fig. S3), with per-base gray wolf ancestry representing 23.2–41.7% (depending on measurement; Table S1). These results are congruent with previous studies that estimated 20–25% from SNP data (vonHoldt et al. 2011), and ∼17–33% using microsatellite data (Roy et al. 1994; Bertorelle and Excoffier 1998). Of note, an anomalous sister-relationship of red wolf to red fox (as outgroup) was supported by 17.9% of the data, a likely result of direct introgression between coyote and gray wolf (Lehman et al. 1991; Gopalakrishnan et al. 2018; Pilot et al. 2019), and/ or inflated discordance due to bottlenecks in contemporary red wolf populations (Brzeski et al. 2014; Waples et al. 2018). Divergence from source genomes was remarkably low, with homozygous ancestry blocks across all chromosomes with D_XY_=0 identified as 49.3% coyote and 54.5% gray wolf.

The distribution of ancestries was notably non-random (Fig. S4-S5), with a substantial enrichment of coyote ancestry on the X-chromosome (Fig. 1A). This was most pronounced in regions of low recombination (<0.5 cM/Mb). It thus comes as no surprise that a significantly higher mean recombination rate was found when gray wolf ancestry blocks were compared between autosomes and X-chromosome (where it is enriched at higher recombination rates; Table 1).

**Table 1:**
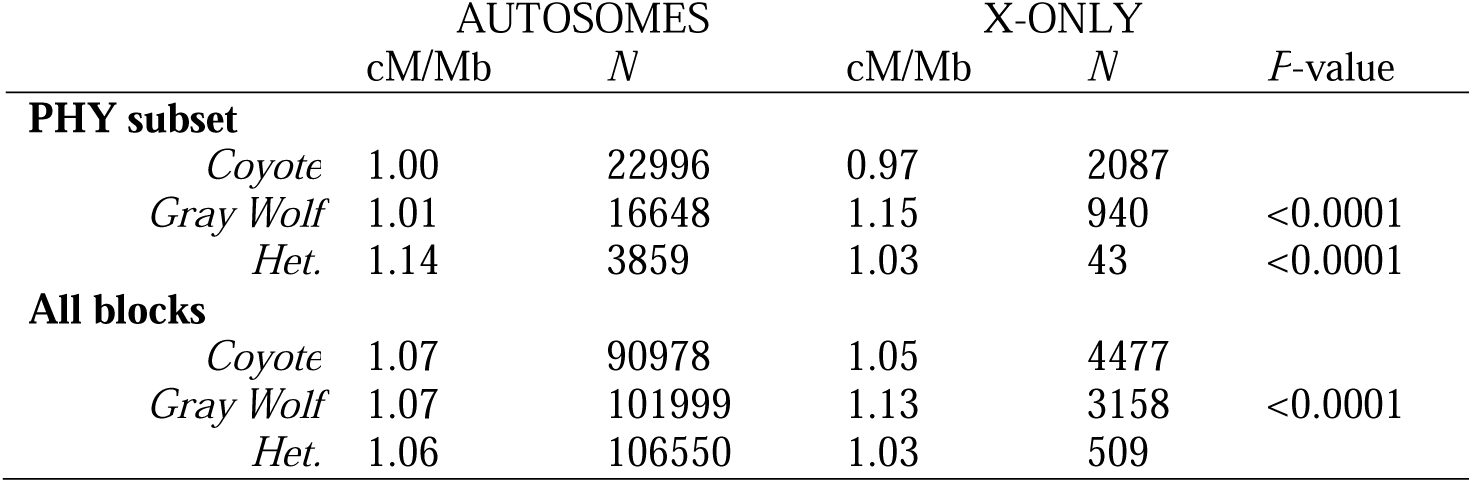
Recombination rate differences between homozygous and heterozygous ancestry blocks in autosomes versus the X-chromosome. Recombination rate is reported as the ratio of centimorgan (cM) per Mb and partitioned into homozygous coyote and gray wolf ancestry and heterozygous ancestry (=HET). Significance is reported for Mann-Whitney *U* test comparing X-chromosome and autosome cM/Mb within each partition.

**Figure 1:**
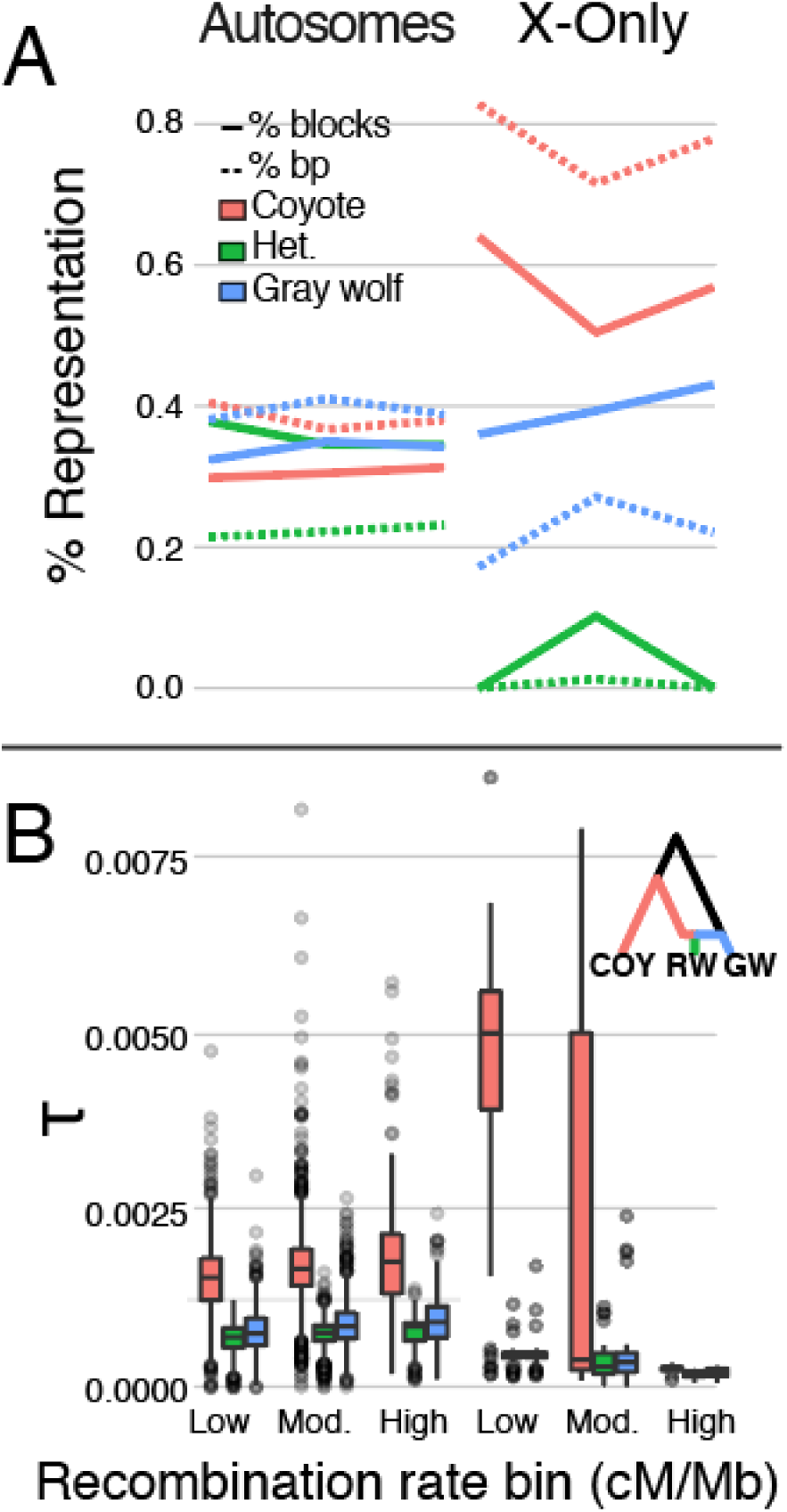
Effect of recombination rate on genomic ancestry proportions (A) and absolute divergence times (B) in the red wolf genome, partitioned among autosomes and X-chromosome. Percent ancestry is computed among genomic ancestry blocks which could be assigned as heterozygous (green), homozygous gray-wolf (blue), or homozygous coyote (red). Genomic representation is reported both as percentage of ancestry blocks (solid) and percentage of base-pairs (bp; dashed). Divergence times (*τ*) measured in expected substitutions were estimated using coalescent hidden Markov models (coalHMM) applied to 1Mb blocks. Results are further partitioned by local recombination rate (cM/Mb) binned as ‘high’ (>=2.0), ‘moderate’ (0.5–2.0), and ‘low’ (<=0.5), and show posterior estimates of: 1) red wolf–coyote divergence (*τ*_COY_; red); 2) red wolf–gray wolf divergence (*τ*_WOLF_; blue); and 3) the time of post-hybridization isolation (*τ*_H_; green).

### Testing the hybrid origin hypothesis

Given the observed distribution of recombination-structured ancestries, two questions emerge with regard to hybridization: (1) Did the temporal context of hybridization contribute to the non-random representation of ancestries? (2) Did hybridization occur as a ‘homoploid hybrid speciation’ event? or (3) Did admixture occur subsequent to a pre-existing isolation? To address these questions, we developed several predictions as a test mechanism in the context of a discriminative framework.

We first recognized the positive relationship between the efficacy of linked selection and the size of linkage blocks in the genome (Nachman and Payseur 2012). Given this, should there be signatures of isolation that pre-date admixture? If so, they would then be expected to occur with highest probability in those regions with low recombination. Likewise, introgressed ancestries are more probable within high-recombination regions where deleterious alleles can be more readily decoupled from neutral or beneficial surroundings (Schumer et al. 2018).

We found positive results in the non-random distribution of ancestries, where a more pronounced occurrence of enriched coyote ancestry was seen within low-recombination regions of the X-chromosome. To expand on this, we also predicted if introgression does indeed mask prior isolation, then those affected genomic regions would display a more shallow coalescence with respect to divergence events (Rosenberg and Feldman 2002; Leache et al. 2014). Juxtaposition of these predictions allowed us to test the hypothesis of hybrid origin versus secondary admixture: If older divergences predominate in areas of low recombination, then the ‘original’ branching pattern is retained (e.g. Fontaine et al. 2015). By partitioning divergence according to recombination rate, we can then unmask ancestral divergence previously obscured.

We fitted a coalescent hidden Markov model (coalHMM) implementing admixture (Cheng and Mailund 2020) to 1Mb blocks of the red wolf genome. We did so to obtain local estimates for red wolf divergence times with regard to coyote (*τ*_COY_) and gray wolf (*τ*_WOLF_) progenitors, as well as putative estimates of post-gene flow isolation (*τ*_H_). In so doing, we uncovered a marked disparity in the range of these estimates between autosomes and the X-chromosome (Fig. 1B and S6–S8). The autosomal estimates were reasonably homogenous across recombination rate bins. This was not so on the X-chromosome: While *τ*_WOLF_ and *τ*_H_ were relatively consistent among recombination rate bins, *τ*_COY_ suggested 20-times older divergence in regions where cM/Mb < 0.5 than in regions where cM/Mb > 2.0 (*μ*=0.004 versus *μ*=0.0002). Thus, divergence was found to be substantially higher in low-recombination regions of the X-chromosome, with younger branching times instead dominating high-recombination regions and the autosomal genome.

We then sampled ∼6000 putatively neutral regions from each parental red wolf sub-genome (following Freedman et al. 2014; vonHoldt et al. 2016) as a means of applying the same demographic modelling approach used in previous studies (i.e. g-PhoCS; Gronau et al. 2011). Results indicated much younger age estimates than those from the COALHMM approach (Fig. 2 and S9), with a mean posterior mutation-scaled divergence time *τ*_COY_=3.8×10^−5^ and *τ*_WOLF_=5.9×10^−6^ (Table S2). Assuming a generation time of three years and an average per-generation mutation rate of 4×10^−9^ (vonHoldt et al. 2016), these correspond to ∼28,500 and ∼4,425 years, respectively. These are congruent with COALHMM estimates taken from high recombination regions of the X chromosome. Interestingly, these results echo a known effect wherein the inclusion of introgressed gene histories promotes ‘tree compression,’ or an underestimation of divergence times (Leache et al. 2014). We also noted several long contiguous blocks showing complete loss of heterozygosity (LOH), in some cases stretching >25Mb (Fig. 3). However, there was no difference in branching time estimates among LOH and non-LOH segments (Fig. 3).

**Figure 2:**
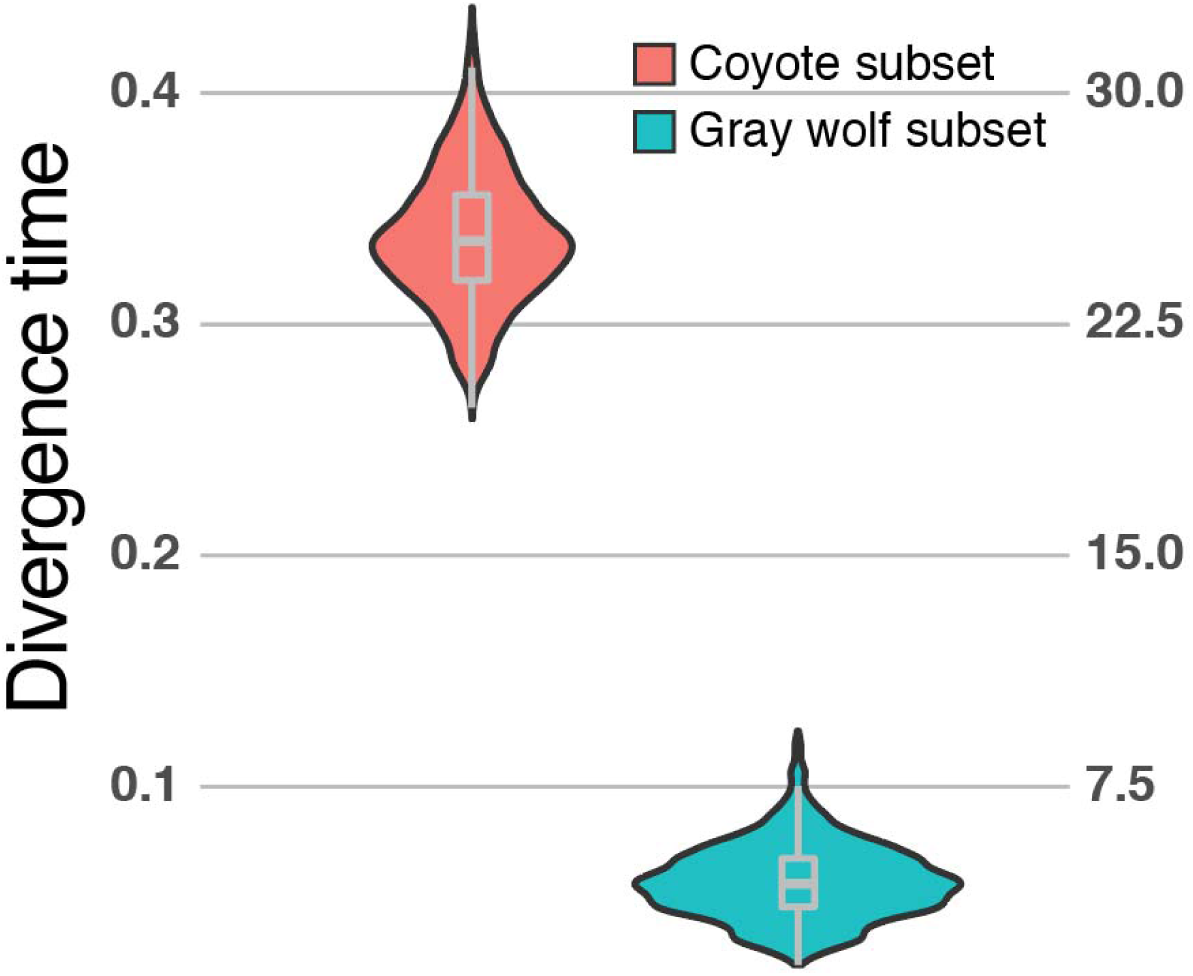
Divergence times estimated using G-PHOCS applied to parental sub-genomes of the red wolf. Divergence times (*τ*) measured in expected substitutions are shown for the red wolf– coyote divergence (*τ*_COY_; red) and red wolf–gray wolf divergence (*τ*_WOLF_; blue). Values are scaled up by a factor of 10,000 (left y-axis) and also provided in calibrated form (right y-axis) in thousands of years, assuming a generation time of three years and an average per-generation mutation rate of *μ*=4×10^−9^ / base pair.

**Figure 3:**
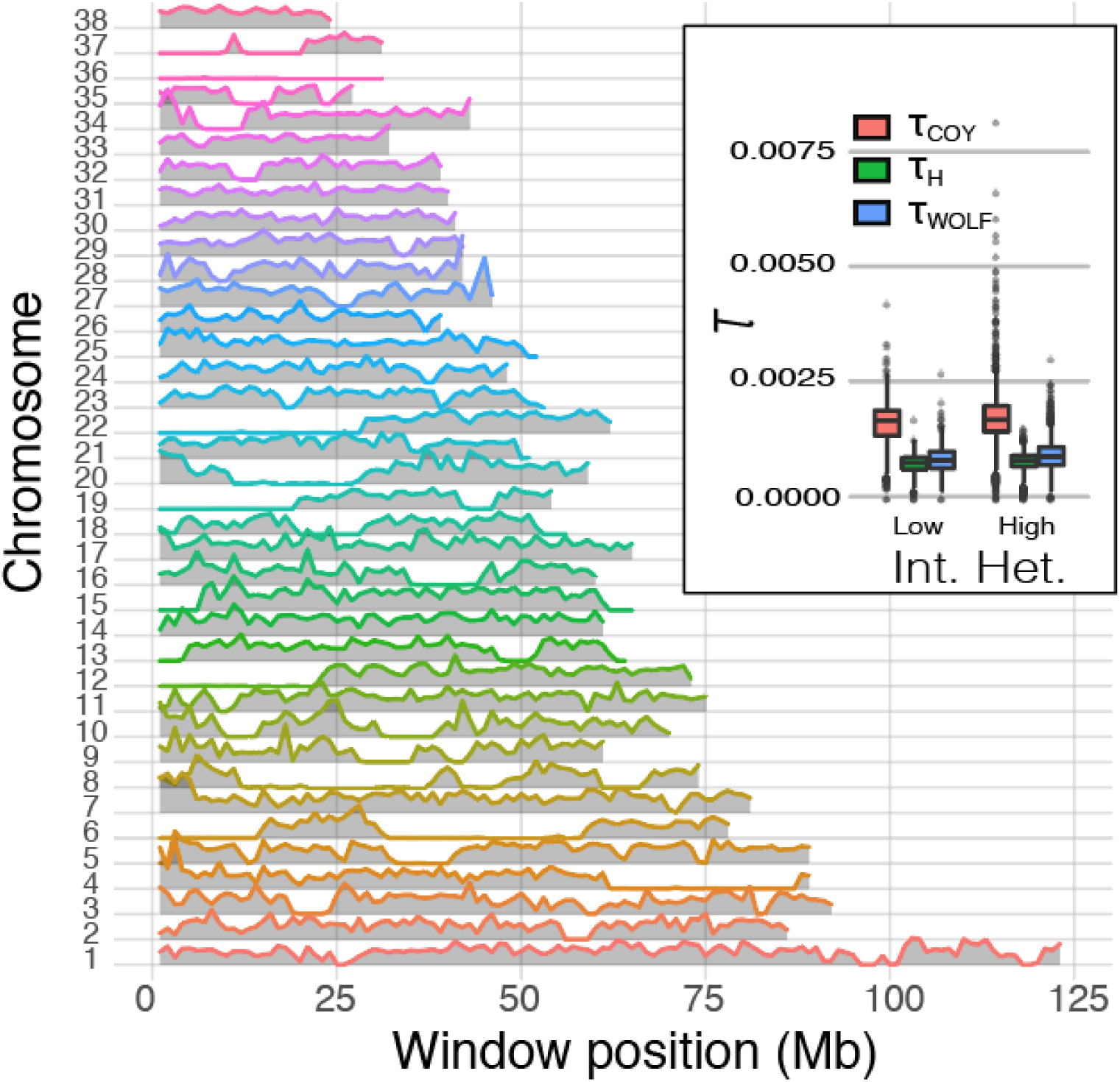
Distribution of interspecific heterozygosity in the red wolf genome, and inferred divergence ages (inset) within low (<0.1 mean interspecific heterozygosity) and high (>=0.1) regions of autosomes. Interspecific heterozygosity was computed as a weighted mean among delimited ancestry blocks encompassed by 1Mb non-overlapping sliding windows.

The observed multi-modality estimates in divergence time suggest multiple separate exchanges between red wolf and putative progenitors (Fig. 1B and S8). To assess this, we took advantage of another prediction: The expected decline in lengths of ancestry blocks over time, as a product of meiotic recombination (Gravel 2012; Ni et al. 2018). Here, the distribution of ancestry-tract lengths in each chromosome was best explained by either two- or three-pulse admixture models (Fig. S10-S11), with the exception of chr9, chr13, and chr37 which fit more appropriately with a gradual admixture model (e.g. with the rate of gene flow continually declining with time since an initial event). The timing of the most recent admixture among those displaying multiple waves (N=36/39) was estimated to be within the last few hundred generations. Older admixtures had a more diffuse distribution, ranging from ∼250–2000 generations (Fig. 4). Mean admixture proportions for distinct pulses ranged from 0.286–0.533 for coyote, and 0.367–0.557 for gray wolf, although the estimated variance in older events was elevated (Fig. S12).

**Figure 4:**
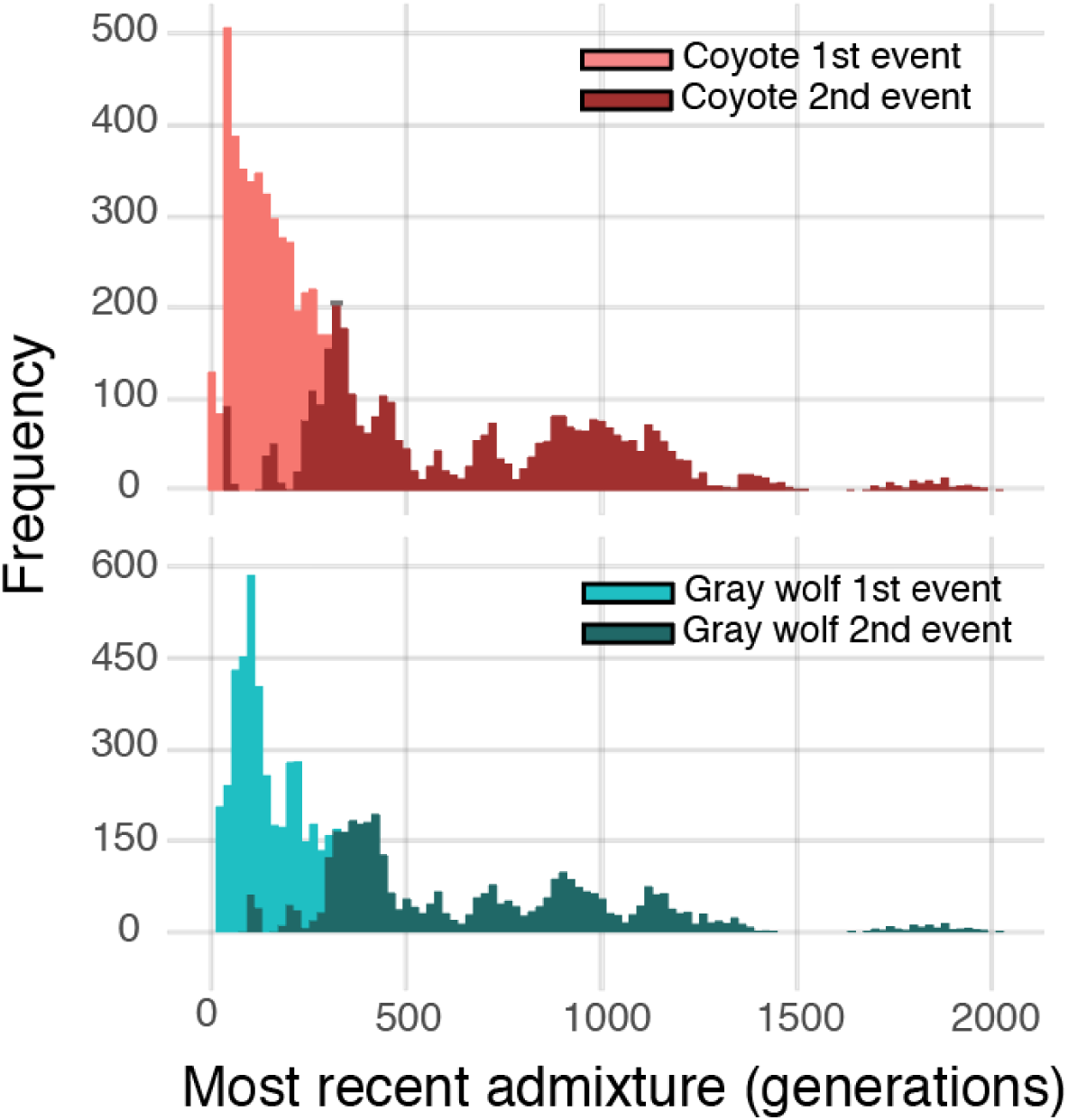
Times of inferred admixture events from coyote (above) and gray wolf (below) into the red wolf genome, measured in generations before the present. Results are shown aggregated from all chromosomes, excluding N=3 chromosomes for which a single-pulse gradual admixture model was selected.

## Discussion

Our findings suggest that extensive secondary introgression, as facilitated by increased permeability of autosomes relative to the X-chromosome, effectively obscured the pre-existing divergence of red wolf. In this sense, the autosomal genome is comparatively homogenous (Fig. 1A), with a low raw divergence and a systematic under-estimation of divergence times stemming from the predominance of introgressed ancestry (Fig. 1B). These results provide quantitative data in support of previous studies that found disproportionate retention of ancient branching patterns in low-recombining regions of sex chromosomes (Fontaine et al. 2015; Schumer et al. 2018; Edelman et al. 2019). This stems from a simultaneous reduction in the rate at which contiguous phylogenetic histories are degraded by linkage, as well as the bolstered efficacy of selection in purging deleterious introgressed elements (Nachman and Payseur 2012; Martin et al. 2019). Moreover, our replication of previous studies (Fig. 2) yielded substantially younger divergence estimates than those from previous studies partitioned by chromosome and recombination rate (Fig. 1B). Our observations agree with prior studies in underscoring the presence of ‘tree compression,’ or a branch lengths reduced/ distorted due to an inability to partition introgressed fron non-introgressed ancestries (Leache et al. 2014; Bangs et al. 2018).

The disparity between autosomes and the X-chromosome reiterates an established phenomenon found in those taxa exhibiting XY and ZW sex determination systems (Fontaine et al. 2015; Seixas et al. 2018; Martin et al. 2019). It is consistent with a ‘large-X effect’ that predicts loci contributing to reproductive isolation accumulate disproportionately on the X- (or Z-) chromosome (Coyne and Orr 1989; Van Belleghem et al. 2018; Presgraves 2018; Runemark et al. 2018). Our results also demonstrated an enrichment of coyote ancestry in low-recombination regions of the X-chromosome, whereas shallower divergence was found within high-recombination regions. A similar logic was presented in Fontaine et al. (2015), wherein a phylogenomic study of *Anopholes* mosquitos also revealed extensive conflict between autosomes and the X-chromosome in the locally dominant branching pattern. They reasoned that gene trees whose branching patterns reflect that of speciation rather than secondary introgression should exhibit deeper coalescence (Fontaine et al. 2015). Such a scenario would similarly explain the patterns of divergence observed in red wolf. Thus, we posit that while masked by secondary introgression in the majority of the genome, lower X-permeability acts as a barrier to exchange, effectively preserving those coalescent patterns established during a more ancient divergence of the red wolf with a coyote-like ancestor.

### Reconciling conflict among genetic and morphological hypotheses

Previous analyses employing these data sparked considerable disagreement, primarily with regards to the timing of gene flow (vonHoldt et al. 2016; Hohenlohe et al. 2017; vonHoldt et al. 2017). The discrepancy between our results and prior studies is two-fold, and stems from: A failure to consider sequential coalescent signal (e.g. as in COALHMM), and the nature of local ancestries structured throughout the genome. Our results are most consistent with an older divergence between coyote and red wolf, an occurrence long obscured by multiple pulses of contemporary admixture (Fig. 4 and S11).

We interpret our results as reconciling the conflict between inferences based on recent molecular work, and those stemming from analyses of modern and historical skeletal remains (e.g. Nowak, 1992). Indeed, the genome does indeed harbor signals of recent and ancient divergences, as established from multiple waves of admixture that successively degraded archaic branching patterns. The most recent admixture event is potentially associated with contemporary anthropogenic change. In this sense, morphological studies could not demonstrate hybridization until the early 1900s, when specimens began trending towards coyote morphologies (Nowak 1979; Nowak 1992; Nowak 2002).

One prevailing question is the status of red wolf prior to modern admixture. Our data suggests its earlier origin, although an absolute estimate is difficult to establish in that effective population sizes, mutation rates, and generation times are all indeterminate (Hohenlohe et al. 2017). Haplotype block lengths suggest admixture as old as ∼1500–2000 generations (Fig. 4), which would place an upper bound extending into the early Holocene, depending on how generation time is defined. Fossil evidence suggests an ecological niche shift in coyote corresponding to megafaunal extinctions at the Pleistocene-Holocene boundary (Meachen and Samuels 2012). Individual body size during the transition period were intermediate between large Pleistocene individuals and more contemporary counterparts that were comparatively diminutive (Meachen et al. 2014). Response of canids to dietary shifts, demographic instability at the glacial-interglacial interface, and wide-spread shuffling of distributions (Pardi and Smith 2016; Loog et al. 2019) may have promoted interspecific contact. We suggest this scenario has plausibility, given the emerging adaptive role for hybridization now commonly evoked in diverse taxa (Lewontin and Birch 2006; Meier et al. 2017; Jones et al. 2018), to include canids (Kays et al. 2010; vonHoldt et al. 2016).

### Conclusion

We employed a fine-scaled, partitioned analysis of genome-wide phylogenetic patterns in red wolf to show that branching patterns reflecting secondary introgression dominate. We also discovered that the presence of introgression varies throughout the genome, as a product of genomic context (e.g. local recombination rates). Because of this, evidence of older divergence was retained by only a fraction of the historically reduced recombination. These findings highlight the difficulties in studying the prevalence of hybridization within the broader Tree of Life, where a sufficiently large numbers of loci can presumably render a singular species history as transparent (Philippe et al. 2011; Hahn and Nakhleh 2016).

However, two biases hinder this approach: (1) The magnitude of signal among loci is clearly disproportionate (Arcila et al. 2017; Shen et al. 2017); and (2) signatures in the genome are deposited by different processes in a heterogeneous manner (as herein). Methods to correct for these biases must explicitly consider the rate of recombination that effectively drives this discrepancy (Payseur and Rieseberg 2016). A failure to do so with regard to red wolf yielded divergence estimates orders of magnitude less than those suggested by fossil evidence (Nowak 1992; Nowak 2002). We reconciled this discrepancy herein by employing estimates based solely on low recombinant regions of the X chromosome. Given this, a failure to partition distinct coalescent histories (e.g. Springer and Gatesy 2018) may result in some phylogenomic studies being interpreted as an artefact of substantial branch length distortion (Leache et al. 2014). The solution is to reconsider the non-random manner by which phylogenetic signal is retained in the genome. This was possible in our red wolf study due to the presence of substantial *a priori* data, to include chromosomal reference assemblies and high-density linkage maps. There are two stumbling blocks to the widespread application of this approach: Resource-limitations and methodological-deficiencies. Both are crucially important if we are to develop a more mature theory of hybridization as a macroevolution process (per Folk et al. 2018).

## Methods

### Read processing, quality filtering, and genotyping

We used previously published genomes for red wolf (=RW; *Canis rufus*), North American gray wolf (=GW; *Canis lupus*), and coyote (=COY; *Canis latrans*), with the red fox (=VUL; *Vulpes vulpes*) serving as an outgroup (vonHoldt et al. 2016a; Kukekova et al. 2018). Paired-end reads were downloaded from the NCBI SRA (SRR7107787; SRR7107783; SRR1518489; SRR5328101-115) and mapped to the domestic dog assembly (CanFam3.1) using BOWTIE2 (Langmead and Salzberg 2012) with sensitive settings, and excluding discordant pairs and unaligned reads. Further processing, sorting, and indexing was performed in SAMTOOLS (Li et al. 2009). PCR duplicates were filtered in PICARD (Broad Institute; broadinstitute.github.io/picard), followed by indel realignment and base quality recalibration in GATK (McKenna et al. 2010; Van der Auwera et al. 2013) as preparation for the HAPLOTYPECALLER pipeline using the ‘Best Practices’ workflow. Genotypes were then inferred jointly using GATK GENOTYPEGVCFS, followed by post-processing, quality filtering, and merging of variant and indel calls.

### Genome-wide phylogenetic patterns

To examine topological and coalescent patterns, we first delimited ancestry blocks within full chromosomal pseudoalignments using a conservative phylogenetic approach, then built pseudoalignments from variant data using a custom Python code (github.com/tkchafin/vcf2msa.py). One issue with this approach is that one cannot assume the genomic reference state for a given nucleotide position will be consistent across the sampled genomes. Thus, within each genome, non-polymorphic bases were treated as un-callable (“N”) when local read depth was < 5. A single-pass algorithm was then used to examine variants (SNPs) for failure of the four-gamete condition (FGT; Hudson and Kaplan 1985). Given the resulting set of incompatible intervals, we then resolved a minimum set of ancestry breakpoints for which no FGT incompatibilities persisted (available as open-source; Chafin 2020).

Delimited blocks were then assigned ancestry using a phylogenetic method. Here, we computed a maximum likelihood estimate (MLE) in IQ-TREE (Nguyen et al. 2014) using integrated model selection and optimization of rate parameters. We discriminated weakly supported relationships by additionally calculating likelihoods under constrained topology searches for each possible quartet resolution, and testing for significant exclusion of alternatives from the MLEs by calculating a bootstrap proportion computed using the RELL approximation (Kishino et al. 1990). Sources of mixed support as a result of systematic errors were differentiated within a given block. For example, the incorrect spanning of recombination events (resulting in concatenated ancestry blocks) was separated from that due to unphased hybrid diplotypes by measuring interspecific heterozygosity. This was derived as the fraction of fixed nucleotide polymorphisms between coyote and gray wolves that were heterozygous for red wolf.

### Testing for multiple-pulse and gradual admixture

Hybrid ancestries are expected to be arranged in large contiguous blocks following an admixture event, with the size of linkage blocks subsequently breaking down over time (Baird et al. 2003). The distribution of ancestry tracts lengths post-admixture can thus be used to understand the timings of genomic contributions (Gravel 2012; Liang and Nielsen 2014), as well as to discriminate multiple-pulse versus continuous admixture models (Zhou et al. 2017; Ni et al. 2018).

To explicitly test among these scenarios, we built a custom SNAKEMAKE pipeline (github.com/tkchafin/multiwaver_snakemake_workflow) for running MULTIWAVER_2.0 (Ni et al. 2018a). To do so, we converted from physical (bp) to genetic (cM) coordinates by utilizing the available comprehensive linkage map for the dog genome (Wong and Neff 2009; Wong et al. 2010). Because the linkage map was built for an earlier version of the assembly (CanFam2), we first converted them using a Python wrapper (available as open-source at github.com/tkchafin/scripts/liftoverCoords.py) for the UCSC LIFTOVER command-line utility (Hinrichs et al. 2006). We then used the LIFTOVER-converted linkage map to construct Marey maps (Siberchicot et al. 2017) for each chromosome, and convert junction positions using cubic interpolation. Ancestries were then assigned to each block based on the phylogenetic results, with blocks having interspecific heterozygosity >0.1 randomly haploidized. We generated 100 independent replicates for each chromosome so as to quantify stochastic variation caused by random ‘pseudo-haploid’ resolution.

### Fitting full-genome admixture histories using coalHMMs

We inferred divergence time parameters using an MCMC (Markov Chain Monte Carlo) sampler for an admixture coalescent HMM (hidden Markov model). HMMs provide a means to probabilistically model transitions along serial or sequential datasets, and are employed widely in genomics and phylogenetics (gene prediction, Stanke and Waack 2003; nucleotide evolution, Yang 1995; Felsenstein and Churchill 1996; and patterns of phylogenetic and geographic diversification, Beaulieu and O’Meara 2016; Caetano et al. 2018). Coalescent HMMs (=coalHMMs) construct a Markov model along a sequence alignment, with ‘hidden’ states as features to reconstruct (Dutheil et al. 2009; Li and Durbin 2011). Hidden states that represent genealogies or coalescent histories are themselves unobservable yet can be predicted from the observed states (=sequence data). Parameters involve processes controlling transitions among hidden states, such as recombination rates (*r*), effective population sizes (*N*_e_), and speciation times (*τ*)(Dutheil et al. 2009). Often (as herein) the primary objective is to infer those demographic parameters from which transition rates are derived (Mailund et al. 2011). In the case of the admixture coalHMM (Cheng and Mailund 2015, 2020), HMMs implementing isolation-with-migration models (Mailund et al. 2012) are combined to generate a pseudolikelihood (or ‘composite’ likelihood) of more complex models that involve multiple lineages. Here we specified priors for the MCMC optimization using demographic estimates from vonHoldt et al. (2016).

Due to the computational complexity of the coalHMM approach (Cheng and Mailund 2015, 2020), the analysis was run separately in 1-million base blocks in two independent replicates per block. We then determined optimal burn-in values using an iterative approach (removing 5% of samples per iteration) using the Geweke diagnostic (Geweke 1992). We also computed effective sample sizes (ESS) for all parameters and assessed convergence of independent chains using the Gelman-Rubin convergence test (Gelman and Rubin 1992; Brooks and Gelman 1998) in the R package CODA (Plummer et al. 2006), removing any blocks for which any parameter-wise ESS fell below 100 or having a Gelman-Rubin statistic <1.01.

### Coalescent demographic modeling

As in vonHoldt et al. (2016), we employed a protocol (Freedman et al. 2014) that targeted a reduced set of putatively neutral loci (1kb in length) for demographic modelling (via G-PHOCS; Gronau et al. 2011). We first excluded regions within a 10kb flanking distance of coding genes (Hoeppner et al. 2014), or conserved non-coding elements (CNEs). The latter were annotated using PHASTCONS scores (Siepel et al. 2005) provided for the *Euarchontoglires* clade, as mapped to the mouse genome (mm9) on UCSC (Freedman et al. 2014). CNEs were then defined as contiguous (over 50bp in length) PHASTCONS scores >0.7 (per Freedman et al. 2014). Interval coordinates for both CNEs and coding genes were converted to the CANFAM3.1 coordinate system (Hinrichs et al. 2006). Our filtered VCF, with reference genome and BED file defining excluded regions, were input to a generalized pipeline (Chafin et al. 2018) that allows for discovery of targeted sub-alignments in genomic datasets. Additional constraints targeted sub-alignments with a maximum proportion of 0.5 uncalled (N) or gap bases. We then subtracted regions from this which were identified as having heterozygous ancestry, and further sampled regions which were at least 100kb apart, truncating regions greater than 5kb in length. Resulting intervals were then extracted as full pseudo-alignments using custom Python code (github.com/tkchafin/vcf2msa.py), with an additional constraint that invariant bases for each species retain the reference base only where >5 reads present; lower-coverage bases were treated as un-callable (“N”). These were then divided into ‘sub-genomes’ by querying dominant phylogenetic ancestry assignments, removing alignments shorter than 500bp, resulting in N=6,100 and 6,255 for gray wolf and coyote sub-genomes (N=12225 loci in total). These served as input for demographic inference in G-PHOCS following the same protocol used in prior studies (Freedman et al. 2014; vonHoldt et al. 2016a).

## Acknowledgements

We thank vonHoldt et al. and Kukekova et al. for making their data available via the NCBI public repository. We are also indebted to students, postdoctorals, and faculty who have contributed thoughtful discussion, assistance with computational resources, or otherwise promoted this work: A. Alverson, M. Bangs, J. Boyko, D. Caetano, D. Chaffin, T. Dowling, J. Pummil, B. Martin, S. Mussmann, A. Tucker, P. Wolinski, and Z. Zbinden. Funding was provided by several generous endowments from the University of Arkansas: The Bruker Professorship in Life Sciences (MRD), and the Twenty-First Century Chair in Global Change Biology (MED). Additional analytical resources were provided by the Arkansas Economic Development Commission (Arkansas Settlement Proceeds Act of 2000) and the Arkansas High Performance Computing Center (AHPCC), and from an NSF-XSEDE Research Allocation (TG-BIO160065) to access the JetStream cloud.

## Data availability

All data for this work were taken from public repositories. Source codes developed in support of this work are available publicly on GitHub (github.com/tkchafin and as cited in-text).

## Supplemental Figures and Tables

**Table S1:**
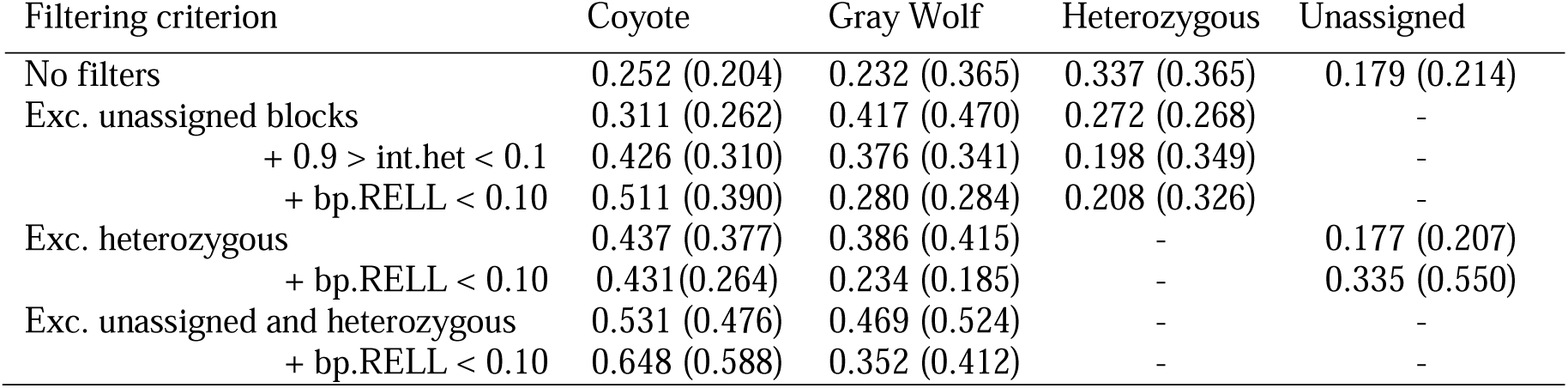
Ancestry proportions of the red wolf genome at varying filtering thresholds and metrics of inclusion. Values are reported as proportion of bases, with proportion of blocks in parentheses. Note that here, ‘Unassigned’ reflects regions in which neither ancestries could be assigned (e.g. red wolf sister to outgroup).

| Filtering criterion | Coyote | Gray Wolf | Heterozygous | Unassigned |
| --- | --- | --- | --- | --- |
| No filters | 0.252 (0.204) | 0.232 (0.365) | 0.337 (0.365) | 0.179 (0.214) |
| Exc. unassigned blocks | 0.311 (0.262) | 0.417 (0.470) | 0.272 (0.268) | - |
| + 0.9 > int.het < 0.1 | 0.426 (0.310) | 0.376 (0.341) | 0.198 (0.349) | - |
| + bp.RELL < 0.10 | 0.511 (0.390) | 0.280 (0.284) | 0.208 (0.326) | - |
| Exc. heterozygous | 0.437 (0.377) | 0.386 (0.415) | - | 0.177 (0.207) |
| + bp.RELL < 0.10 | 0.431(0.264) | 0.234 (0.185) | - | 0.335 (0.550) |
| Exc. unassigned and heterozygous | 0.531 (0.476) | 0.469 (0.524) | - | - |
| + bp.RELL < 0.10 | 0.648 (0.588) | 0.352 (0.412) | - | - |

**Table S2:** Mutation-scaled and absolute divergence time estimates from g-PhoCS. Parameters are as follows: population divergence time for red wolf and coyote (*τ*_COY_); divergence time for red wolf and gray wolf (*τ*_WOLF_); divergence time for all *Canis* species (*τ*_CANIS_); and the divergence time for the root (*τ*_ALL_). Values shown are the raw arithmetic mean estimates, with calibrated estimates in years in parentheses, assuming a generation time of three years and an average per-generation mutation rate of *μ*=4×10^−9^ / base pair.

| Analysis | $\tau_{\text{COY}}$ | $\tau_{\text{WOLF}}$ | $\tau_{\text{CANIS}}$ | $\tau_{\text{ALL}}$ |
| --- | --- | --- | --- | --- |
| Coyote subset | $3.8 \times 10^{-5}$<br>(28,500) | - | $7.3 \times 10^{-4}$<br>(547,500) | $2.5 \times 10^{-3}$<br>(1,875,000) |
| Gray wolf subset | - | $5.9 \times 10^{-6}$ (4,425) | $7.6 \times 10^{-4}$<br>(570,000) | $2.9 \times 10^{-3}$<br>(2,175,000) |

**Figure S1:**
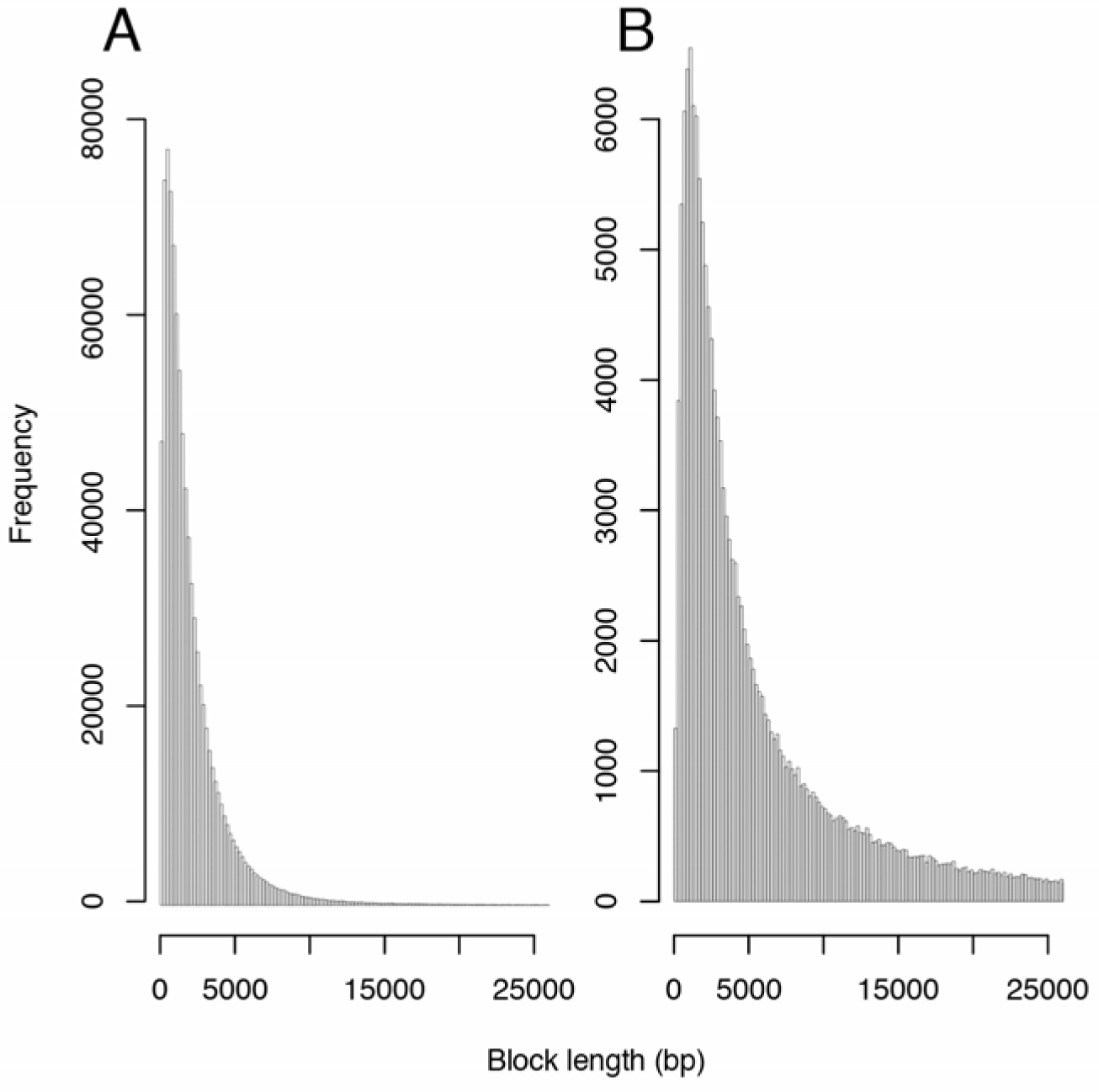
Ancestry block lengths in the red wolf genome before (A) and after (B) merging consecutive blocks of the same ancestry. Note truncation of the x-axis for interpretability.

**Figure S2:**
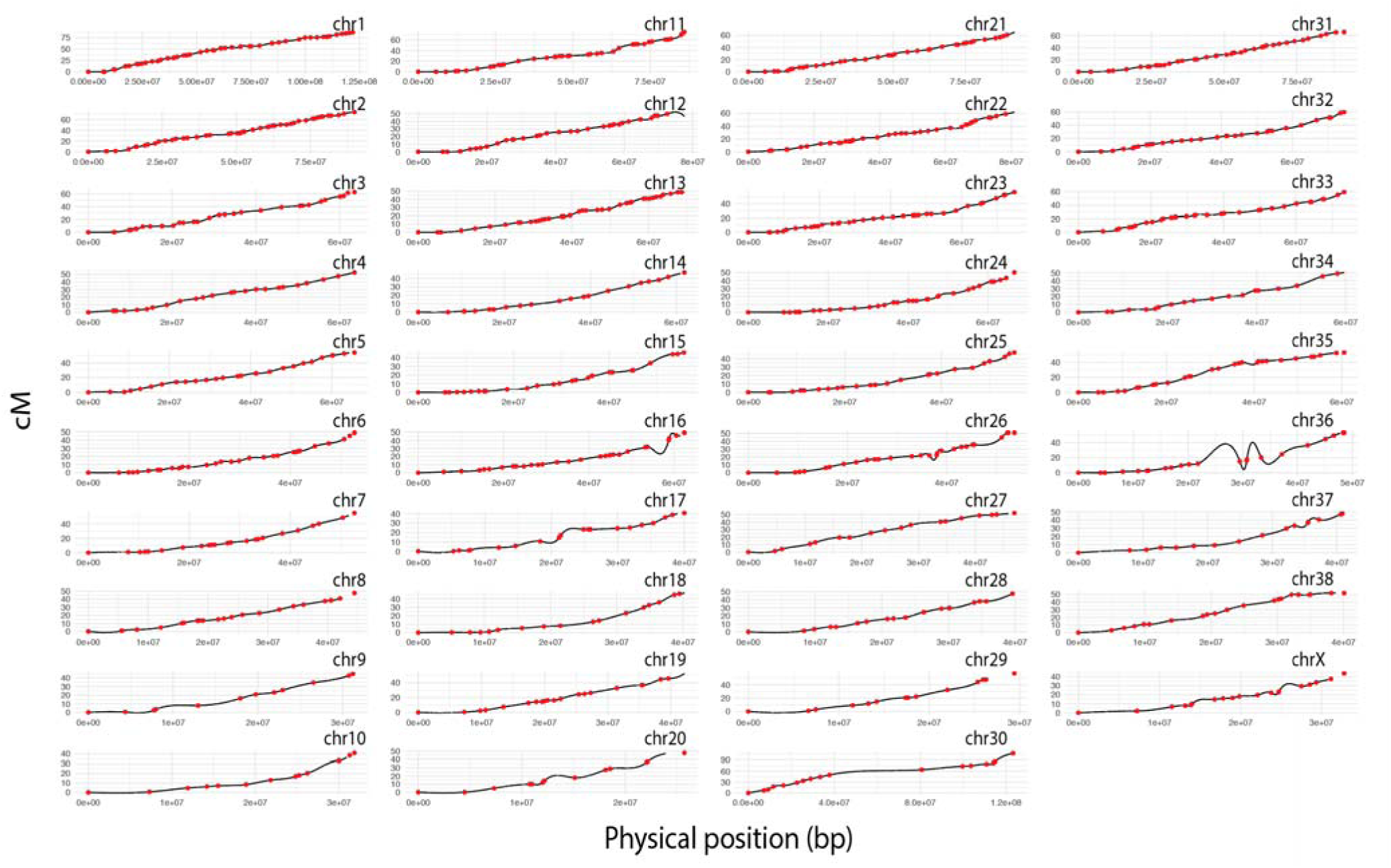
Cubic interpolation models for red wolf chromosomes (black) with points depicting datapoints from the genetic map (red)

**Figure S3:**
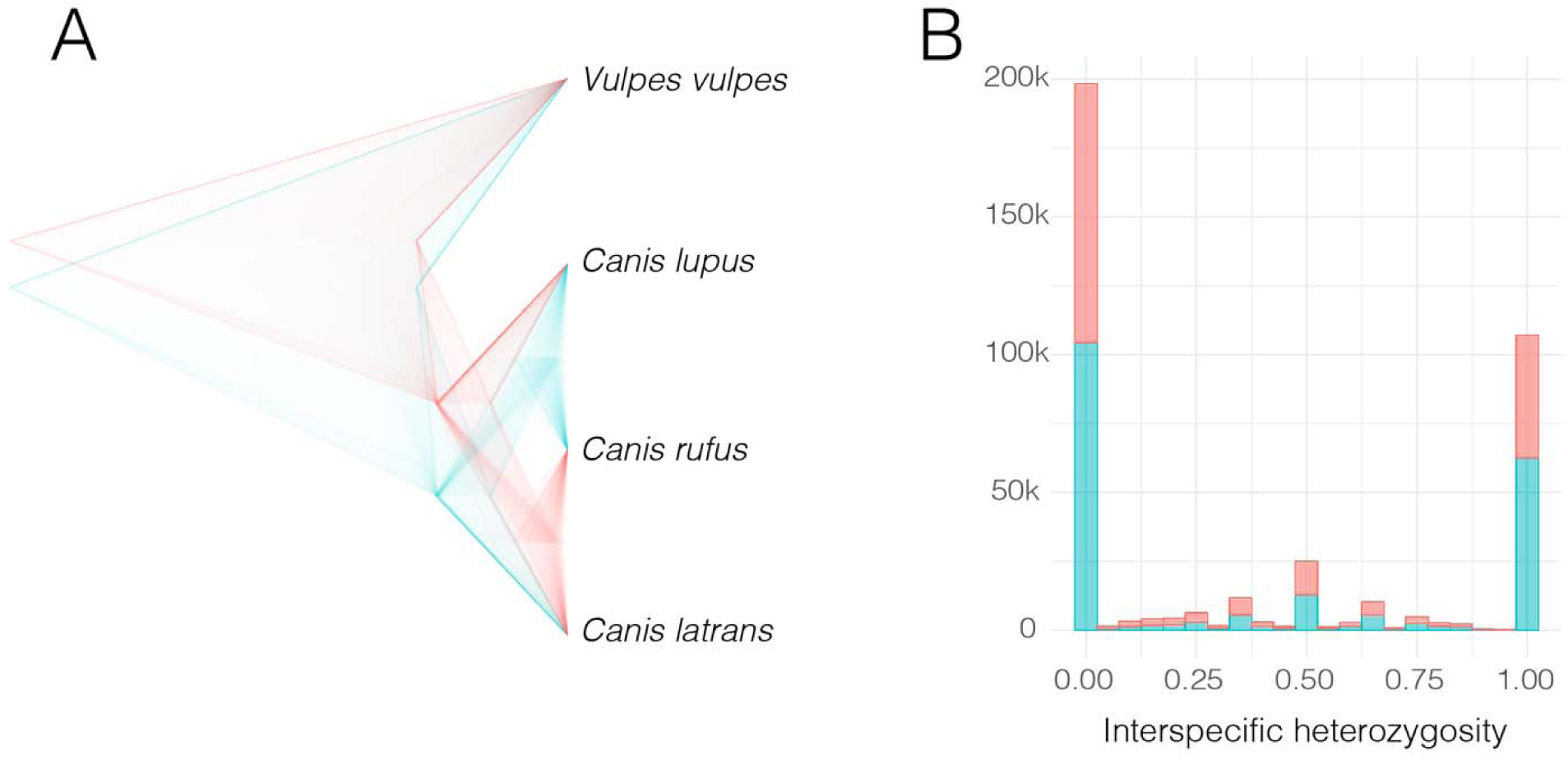
Densitree plot (A) of gene trees and distribution of interspecific heterozygosity (B) among sampled genomic ancestry blocks in the red wolf genome. Gene trees are restricted to those which were significantly supported by approximated bootstrap proportions (e.g. <10% of trees supporting an alternate topology), whereas interspecific heterozygosity is reported for all blocks.

**Figure S4:**
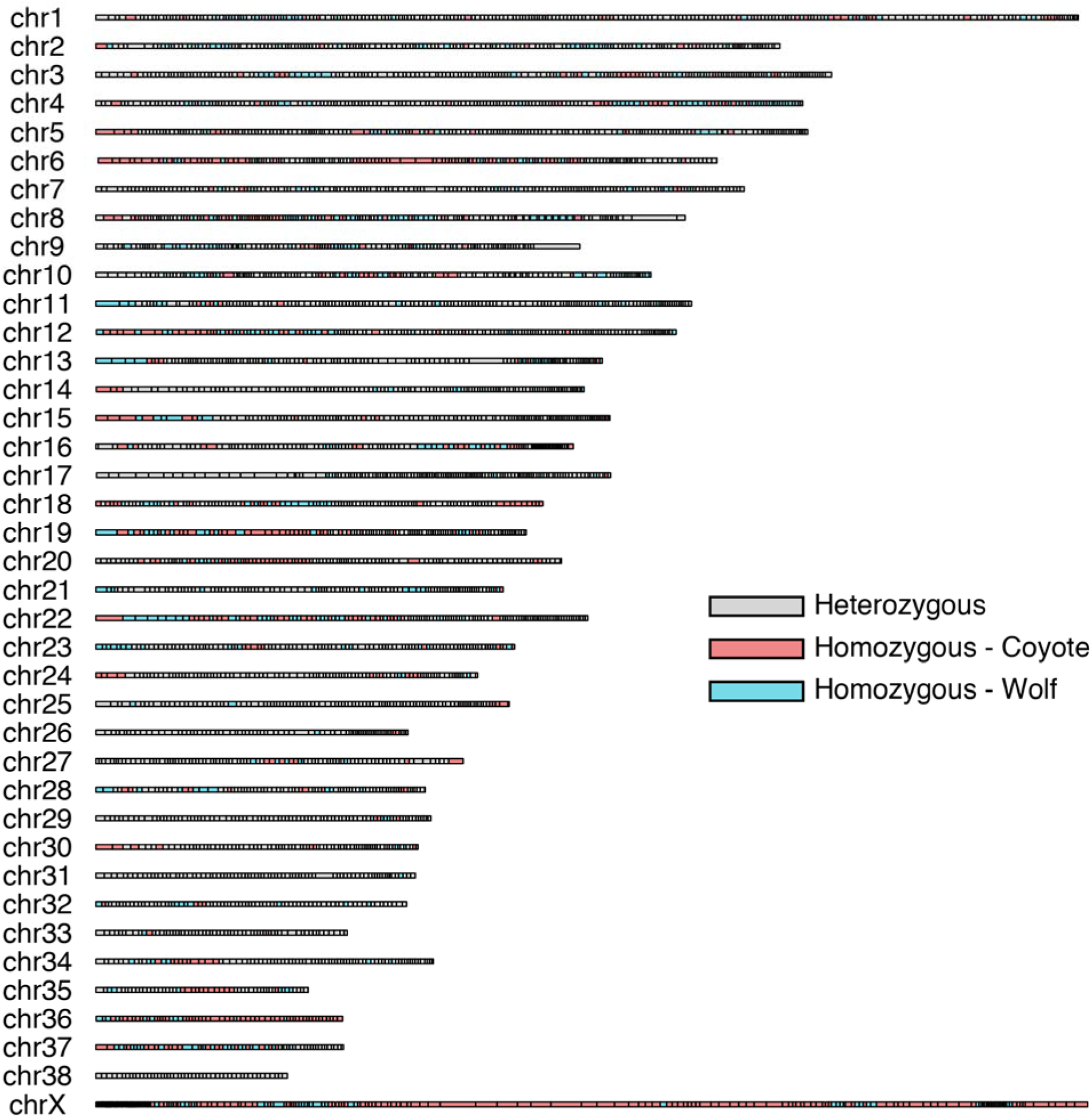
Dominant ancestries summarized along red wolf chromosomes, as determined via ‘majority-rule’ among delimited ancestry blocks merged into 500kb segments, excluding blocks for which ancestry could not be decisively determined.

**Figure S5:**
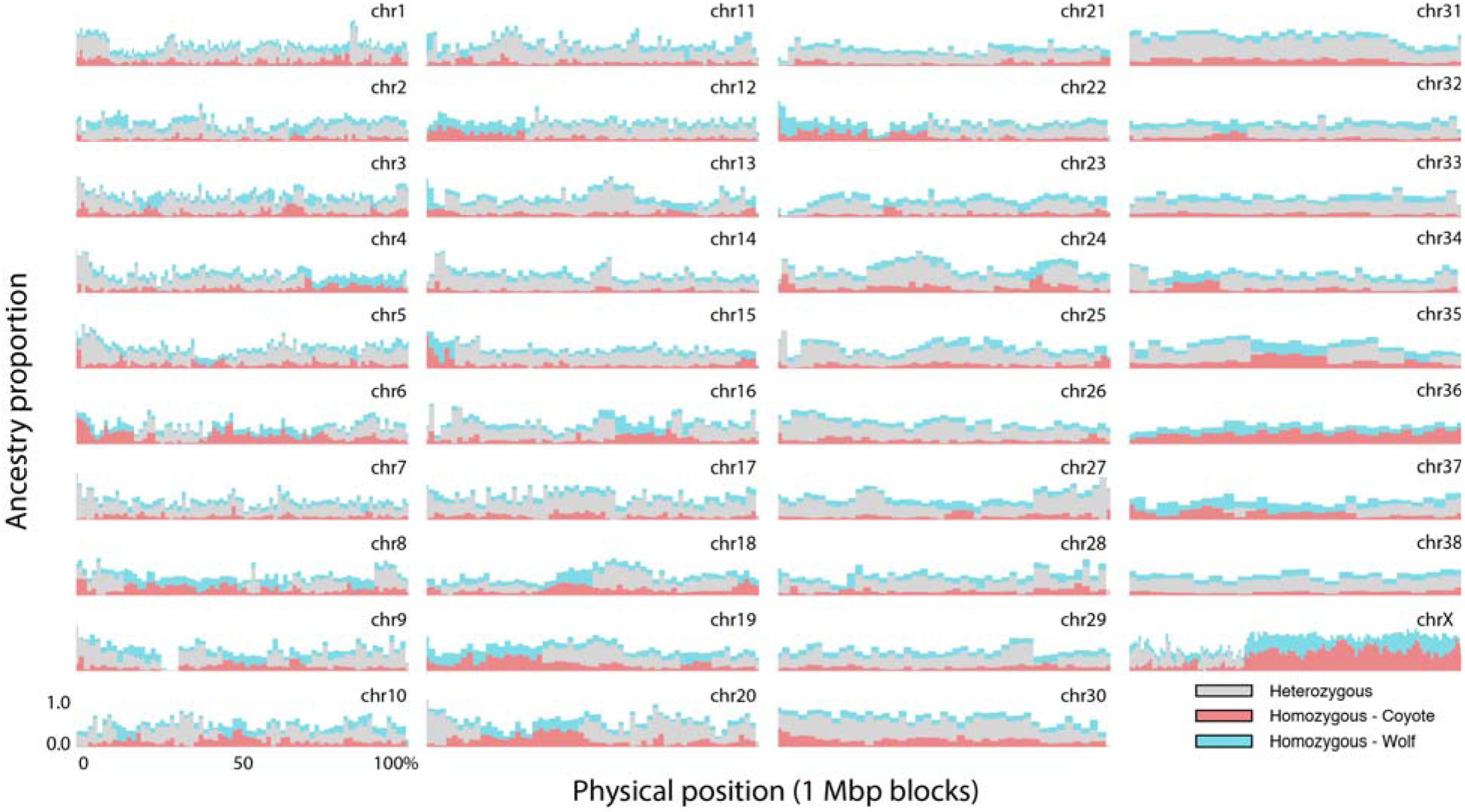
Proportion of assigned ancestries in the red wolf genome in 1 megabase blocks per chromosome, with blocks that could not be conclusively called as heterozygous or parental-homozygous excluded.

**Figure S6:**
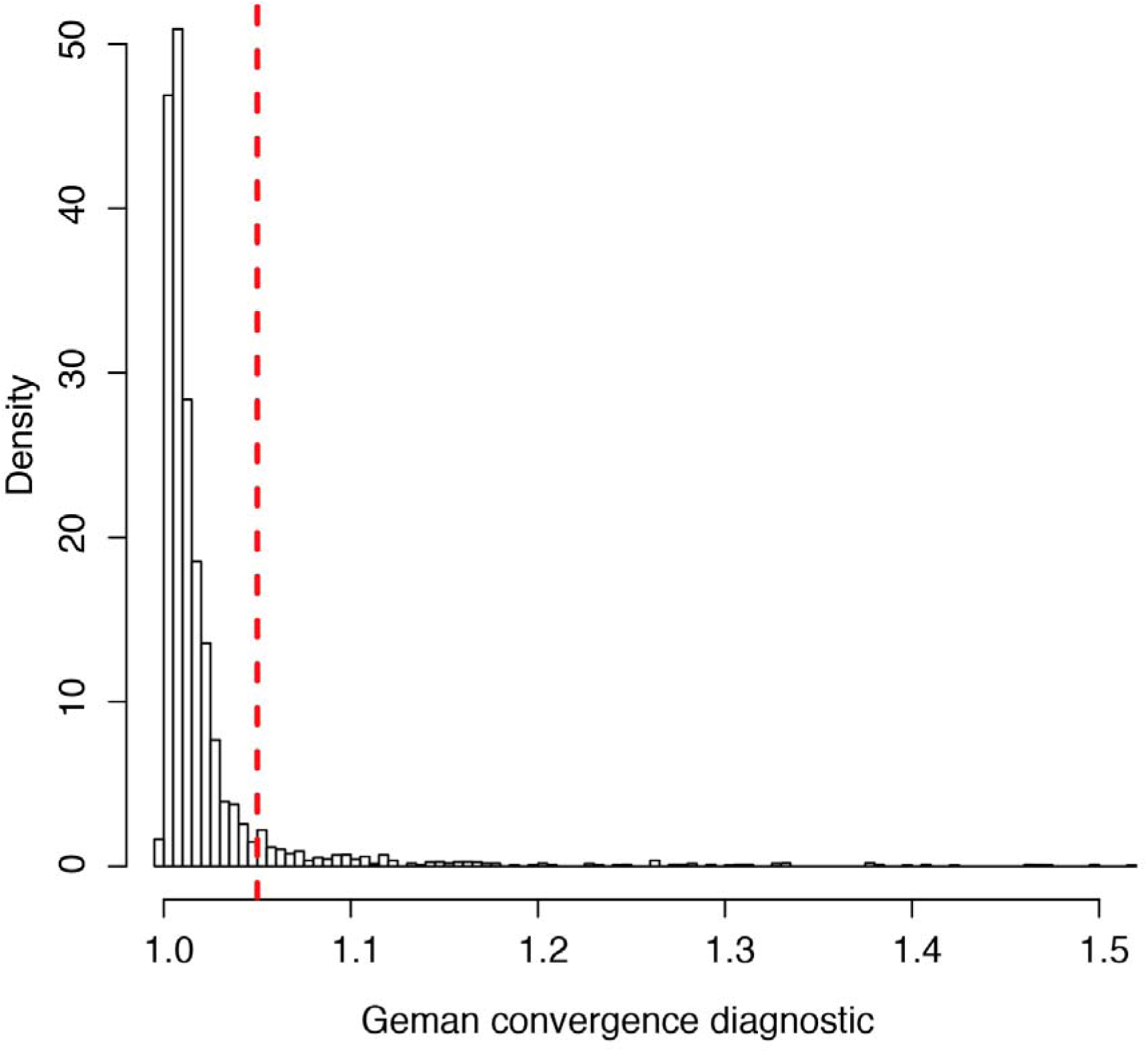
Histogram of Gelman convergence diagnostics across MCMC chains (each having two replicates), showing a cutoff threshold (red) of 1.05. Values shown are post burn-in, following an automated iterative procedure testing burn-in values according to the Geweke diagnostic.

**Figure S7:**
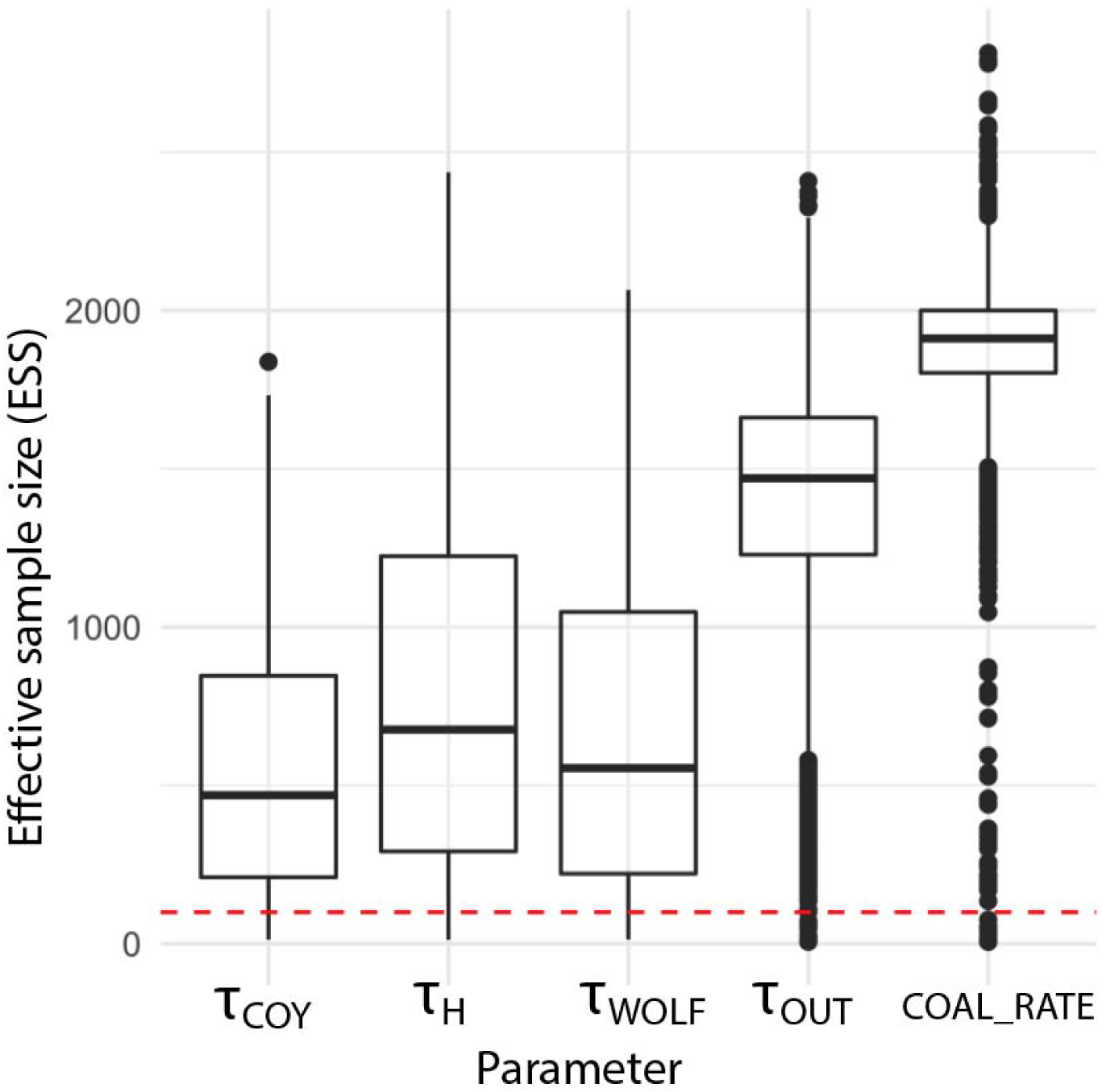
Effective sample sizes summarized across coalHMM MCMC chains passing Gelman-Rubin convergence threshold of 1.05. The minimum ESS threshold (red) of 100 was used to filter coalHMM results.

**Figure S8:**
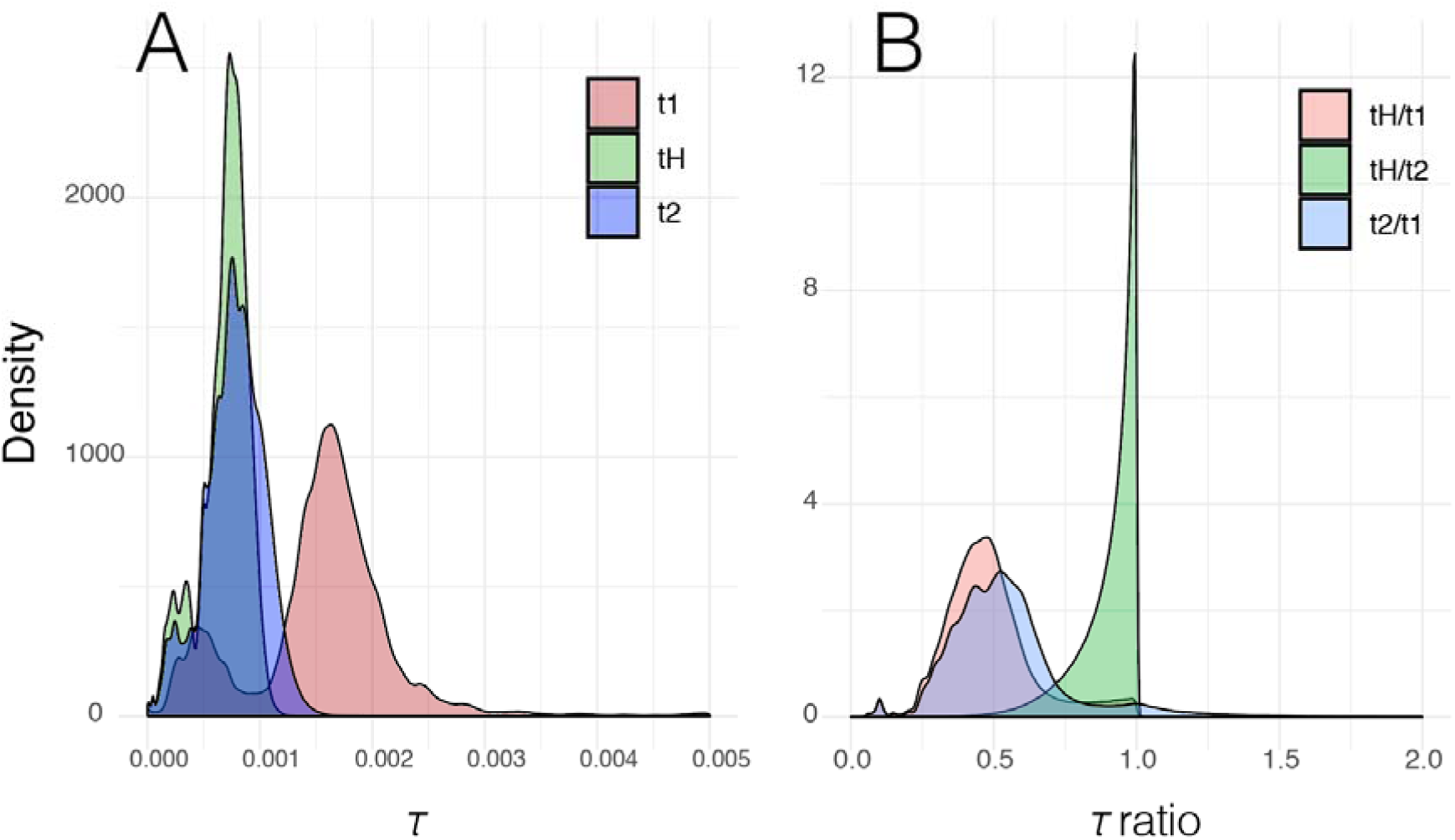
Posterior distributions of coalescent times inferred using coalHMM within 1Mb blocks of the red wolf genome (A) and the ratios among dates (B)

**Figure S9:**
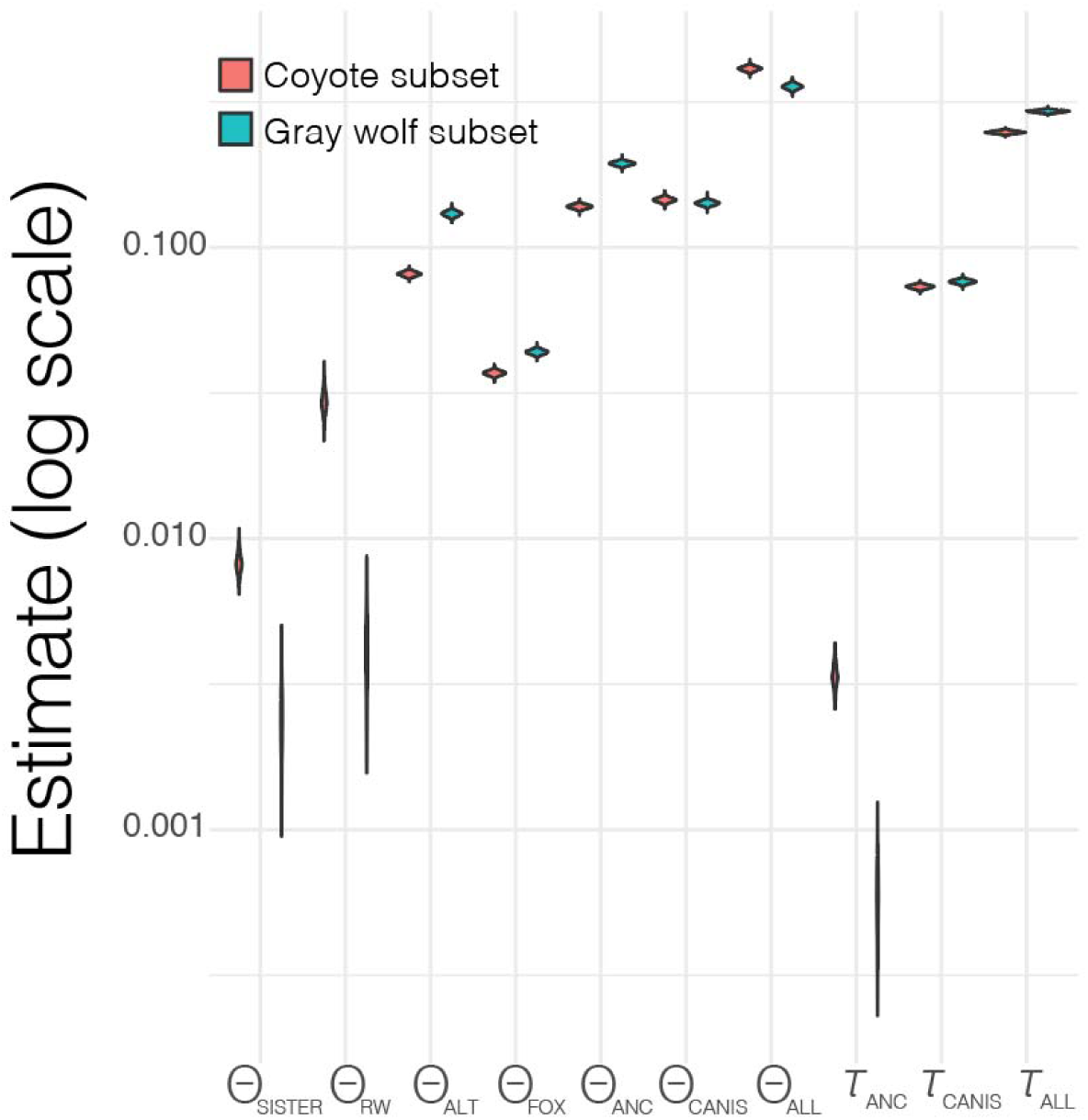
Full set of parameter estimates from g-PhoCS sub-genome partitioned analysis. Results are grouped by putative coyote and gray wolf sub-genomes, showing mutation-scaled effective population size (Θ) and divergence times (*τ*). Both analyses assumed a pectinate tree, wherein the ‘source’ (=parental) genome is sister to the red wolf, with *Vulpes vulpes* as an outgroup. Parameters are displayed on a log scale, and are as follows: Effective population size for the source genome for each respective subset (Θ_SISTER_); effective population size for red wolf (Θ_RW_); population size of the non-source genome (e.g. coyote for gray wolf sub-genome analysis; Θ_ALT_); ancestral population size for red wolf and the source genome (Θ_ANC_); ancestral population size for all *Canis* species (Θ_CANIS_); ancestral population size for the root (Θ_ALL_); population divergence time for red wolf and the source genome (*τ*_ANC_); divergence time for all *Canis* species (*τ*_CANIS_); and the divergence time for the root (*τ*_ALL_).

**Figure S10:**
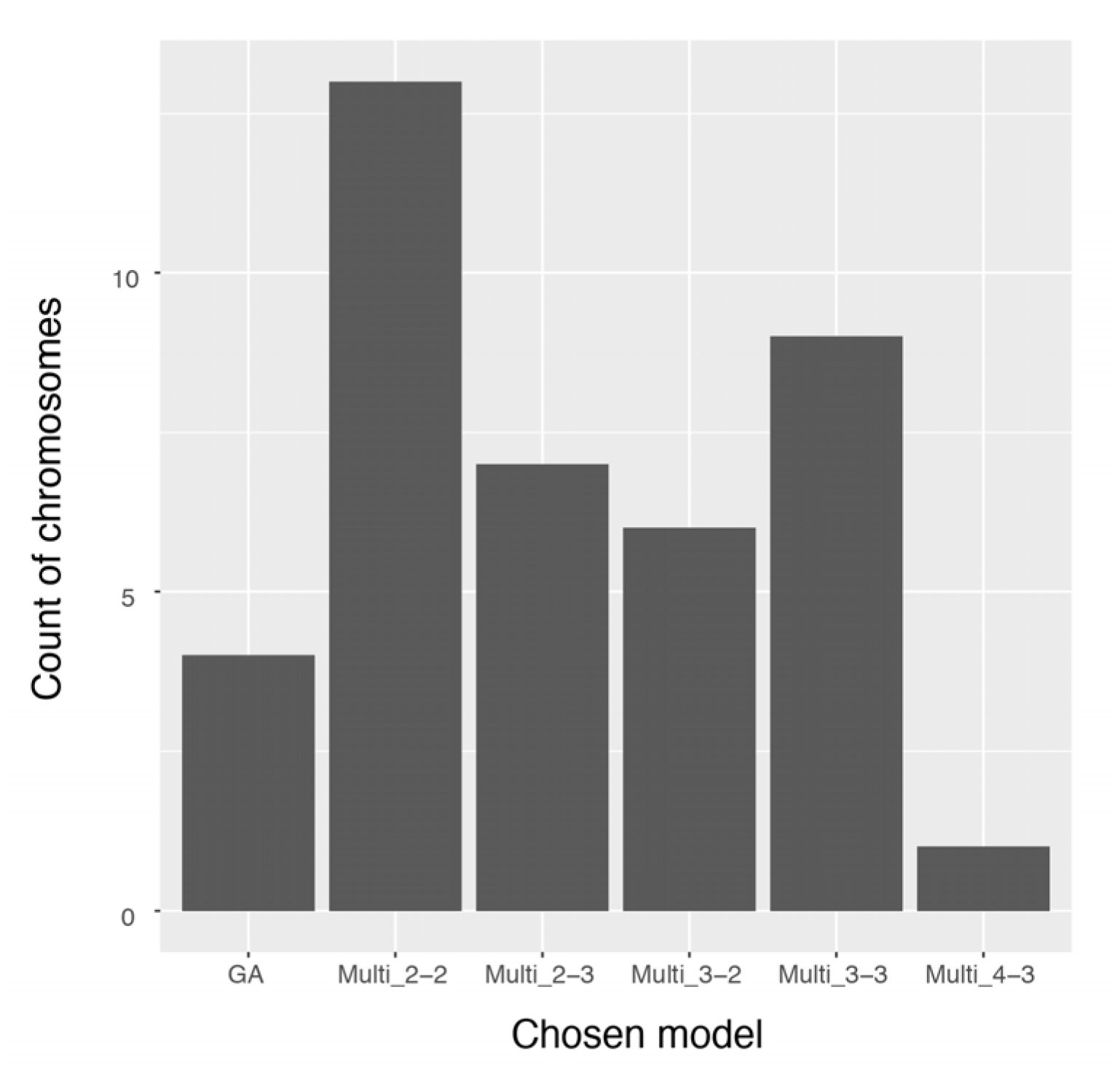
Total count of chromosomes choosing categories of admixture models, where GA=gradual admixture; and Multi_*x-y* represents multiple-pulse admixture models where *x*=number of inferred coyote admixture events and *y*=number of gray wolf events.

**Figure S11:**
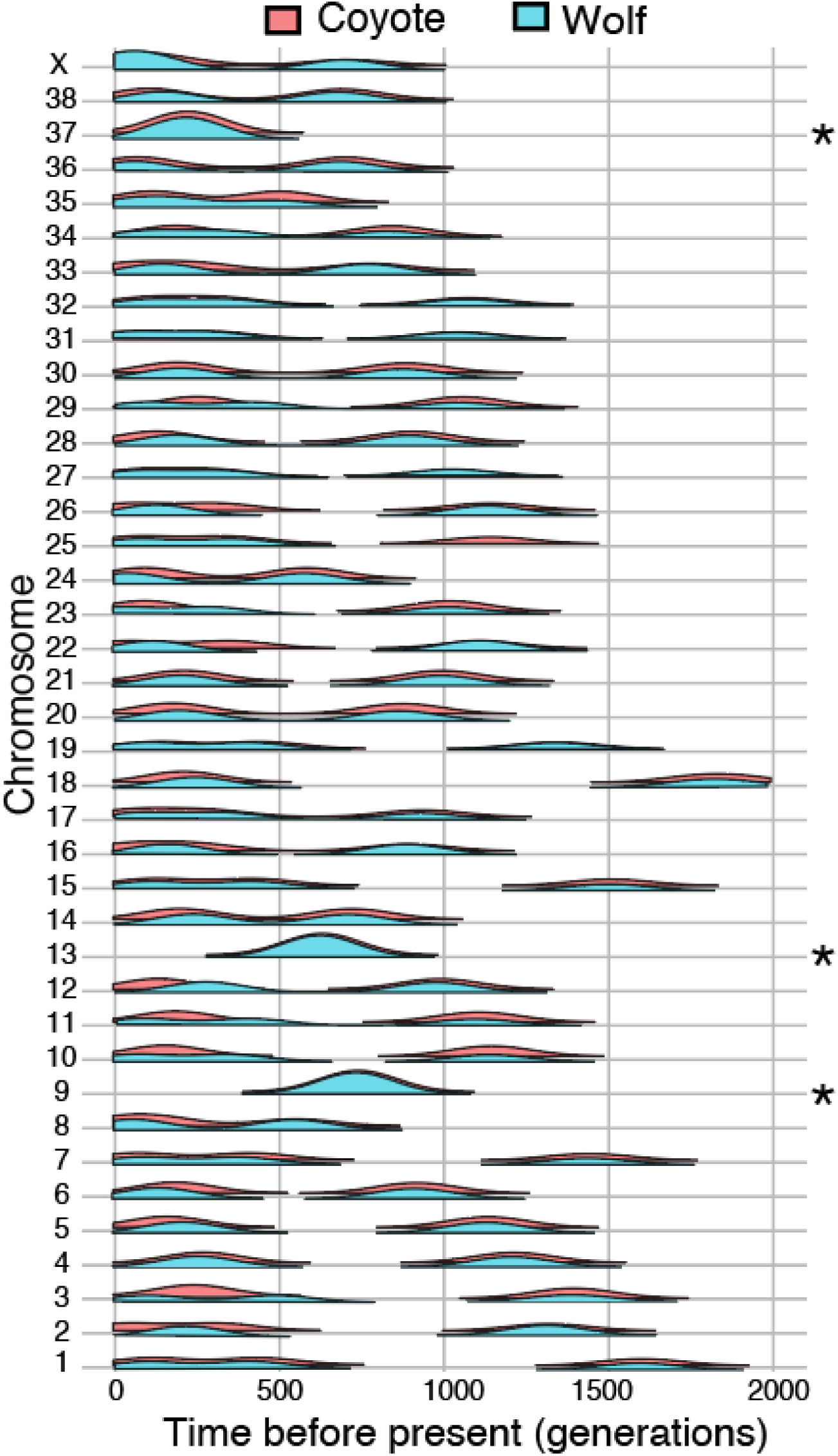
Admixture times inferred from ancestry block lengths in a multiple-pulse hybridization model. Time is shown in generations before the present, with the *y*-axis depicting densities based on 100 replicate datasets per chromosome, wherein heterozygous blocks were randomly assigned ancestry. Note that chromosomes best fitting a single-pulse model (*) depict the inferred start time for a gradual admixture (GA) scenario in which migration rates thereafter towards the present.

**Figure S12:**
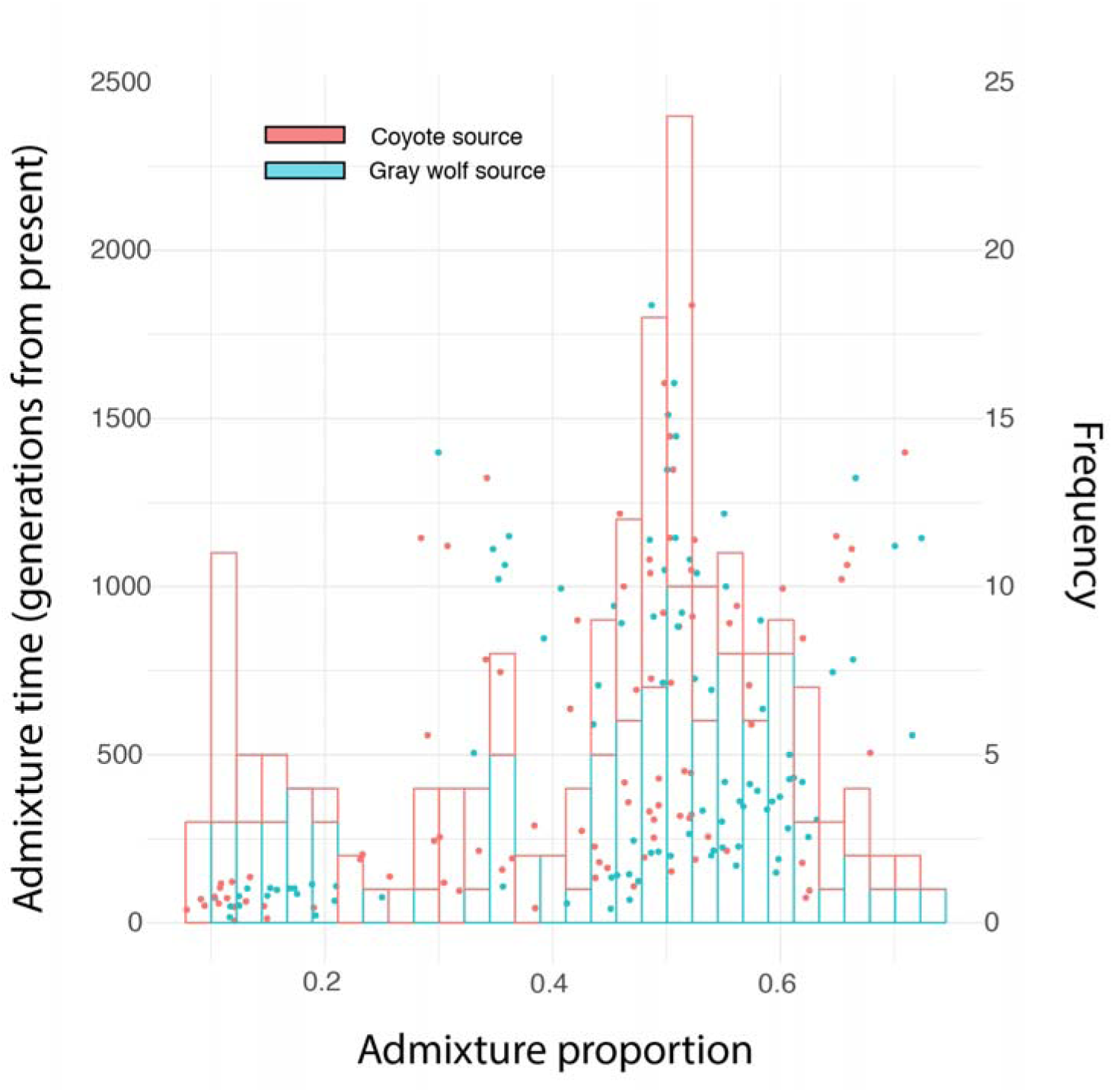
Admixture proportion for inferred admixture events inferred at different times (measured in generations before the present). The left axis (=points) show the measured admixture times as a function of admixture proportions, which the right axis (=histogram) shows frequency of admixture proportions across all events.

